# A distinct inner nuclear membrane proteome in *Saccharomyces cerevisiae* gametes

**DOI:** 10.1101/2021.08.02.454801

**Authors:** Shary N. Shelton, Sarah E. Smith, Jay R. Unruh, Sue L. Jaspersen

**Affiliations:** Stowers Institute for Medical Research, Kansas City, Missouri 64110; Department of Molecular and Integrative Physiology, University of Kansas Medical Center, Kansas City, Kansas 66160

**Keywords:** budding yeast, INM, reticulon, Heh1/Heh2, sporulation

## Abstract

The inner nuclear membrane (INM) proteome regulates gene expression, chromatin organization, and nuclear transport, however, it is poorly understood how changes in INM protein composition contribute to developmentally regulated processes, such as gametogenesis. Using a split-GFP complementation system, we compared the distribution of all C-terminally tagged transmembrane proteins in *Saccharomyces cerevisiae* in gametes to that of mitotic cells. Gametes contain a distinct INM proteome needed to complete gamete formation, including expression of genes linked to cell wall biosynthesis, lipid biosynthetic and metabolic pathways, protein degradation and unknown functions. Based on the inheritance pattern, INM components are made *de novo* in the gametes. Whereas mitotic cells show a strong preference for proteins with small extraluminal domains, gametes do not exhibit this size preference likely due to the changes in the nuclear permeability barrier during gametogenesis.

## Introduction

The double lipid bilayer that forms the nuclear envelope (NE) is composed of the outer nuclear membrane (ONM) and the inner nuclear membrane (INM). The INM and ONM are contiguous at multiple sites that contain nuclear pore complexes (NPCs). These multi-subunit machines control passage of macromolecules into and out of the nucleus, including components of the INM. Transport of proteins to the INM results in a distinct INM proteome in contrast to the ONM, which is contiguous with the ER (Florens et al. 2008; Mudumbi et al. 2016; Schirmer et al. 2003, 2005). Many, if not most, of the unique properties of the NE can be attributed to the INM, for example, its distinct lipid composition, its mechanical rigidity, and its role in chromosome organization. During development, differentiation, and in several diseased states, the NE undergoes several changes in its structure and composition (Chen et al. 2014; D’Angelo et al. 2009; Dauer and Worman 2009; Davidson and Lammerding 2014; Gordon et al. 2014; Grima et al. 2017; Wilkie et al. 2011). Expanding our insight into the protein composition of the INM in different cell types is necessary to facilitate our understanding of pathological processes related to defects in the NE, and more specifically, the INM.

The tightly regulated developmental process of gametogenesis is where a progenitor cell undergoes two consecutive rounds of nuclear divisions, meiosis I and meiosis II, to form four haploid gametes. In yeast, several changes occur at the meiosis I to meiosis II transition, including detachment of mitochondria and endoplasmic reticulum from the cell cortex followed by relocalization to the ONM (Gorsich and Shaw 2004; Miyakawa et al. 1984; Sawyer et al. 2019; Suda et al. 2007). Plasma membrane synthesis at the cytoplasmic side of the spindle pole bodies forms the prospore membrane (PSM). The growing PSM encapsulates nascent nuclei while also capturing a small fraction of the cytoplasm before it closes at the end of meiosis II (Fuchs and Loidl 2004; Knop and Strasser 2000; Moens and Rapport 1971; Neiman 1998). Unexpectedly, some nuclear material such as components of NPCs and the nucleolus are not incorporated into gametes by the PSMs but are left behind in a “fifth compartment” (Fuchs and Loidl 2004). Following meiosis II, vacuolar lysis removes debris, including the fifth compartment, that has been sequestered during gametogenesis before complete gamete maturation (King et al. 2019; King and Unal 2020). The observation that most nucleoporins, with the exception of those bound to chromatin, are left behind in the degraded NE-bound fifth compartment (King et al. 2019), led to the proposal that nucleoporins are either made *de novo* or return from a pre-existing sequestered pool of NPCs in gametes (King and Unal 2020). It is unknown if other NE components are removed from the NE like NPCs or if they are maintained due to their connection with chromatin and/or roles in NE synthesis and structure.

## Materials and Methods

### Library construction

The generation of the C-terminally tagged library in the BY strain, SLJ7859, (*MATα can1Δ:: STE2pr-SpHIS5 lyp1Δ his3Δ1 leu2Δ0 ura3Δ0 met15Δ0 LYS2 pCEN/ARS-LEU2*-*NOP1pr-GFP*_*11*_*-mCherry-PUS1* (pSJ1321)) has been described previously (Smoyer et al. 2016). Briefly, genes associated with the GO annotation of integral component of membrane or transmembrane were compiled using the *Saccharomyces* Genome Database (SGD); TMHMM (Krogh et al., 2001) was used to predict additional genes containing hydrophobic stretches of greater than 16 amino acids using a version of the genome downloaded on June 10, 2012. GFP_1-10_ was integrated using PCR-based tagging at the C-terminus of those identified target genes in the haploid strain. Correct integration was verified by PCR.

### INM gamete screen

The BY4742 derivative (*MATα can1Δ:: STE2pr-SpHIS5 lyp1Δ his3Δ1 leu2Δ0 ura3Δ0 met15Δ0 LYS2 pCEN/ARS-LEU2*-*NOP1pr-GFP*_*11*_*-mCherry-PUS1*) haploid strain containing the C-terminal GFP_1-10_ tagged genes was mated to the SK1 (MATa ho::*LYS2 lys2 ura3 leu2*::hisG *his3*::hisG *trp1*::hisG *leu*::*Nop1pr*-*GFP*_*11*_*-mCherry-PUS1*-*LEU2)* haploid strain using the Singer ROTOR (Singer Instruments) on YPD plates for 2 d at 30°C. Diploid hybrid strains then went through two cycles of growth on SD-Leu for 3 d at 30°C before making glycerol stocks.

To screen for INM access, diploid hybrid strains were grown for 24 h at 30°C in enriched synthetic complete media (eSCM) lacking leucine (eSC-Leu: 4% glucose, 0.67% bacto-yeast nitrogen base without amino acids, 0.2% Leu amino acid dropout powder) with agitation. Following 24 h of growth, cells were washed twice with water and twice with sporulation (SPO) media (1% potassium acetate, 0.32% of amino acid dropout powder without methionine, 0.2% methionine). Cells were then resuspended in SPO and shaken at 30°C for 48 h. To facilitate screening in live cells, cell growth was staggered.

An aliquot of cells was immobilized between a glass slide and a no. 1.5 coverslip before imaging with a Perkin Elmer (Waltham, MA, USA) Ultraview spinning disk confocal microscope equipped with a Hamamatsu (Hamamatsu, Japan) EMCCD (C9100-13) optimized for speed, sensitivity and resolution. The microscope base was a Carl Zeiss (Jena, Germany) Axio-observer equipped with an αPlan-Apochromat 100x 1.46NA oil immersion objective and a multiband dichroic reflecting 488 and 561 nm laser lines. GFP images were acquired with 488 nm excitation and 500-550 nm emission. mCherry images were acquired with 561 nm excitation and 580-650 nm emission. Data were acquired using the Perkin Elmer Velocity software with a z spacing of 0.4 *µ*m. Exposure time, laser power and camera gain were maintained at a constant level chosen to provide high signal-to-noise but avoid signal saturation for all samples. Images were processed using Image J (NIH, Bethesda, MD). Maximum-intensity projections over two to five *z*-slices were created, and images were then binned 2×2 with bilinear interpolation. A complete repository of all processed images is available at https://research.stowers.org/imagejplugins/jaspersen_meiosis_splitGFP_screen/.

Localization to the INM, nucleus or other compartments was determined by manual inspection of the images and is summarized in Table S1. Bioinformatic analysis of hits in this screen and a previous published screen of the mitotic INM (Smoyer et al. 2016) was performed using Yeast Mine tools at SGD. The size and topology were previously determined using Phobius and TMHMM (Smoyer et al. 2016). Protein levels of hits from the screen were extracted from previously published data (Cheng et al. 2018) and normalized so that protein expression values throughout meiosis fall between 0 and 1.

### Yeast strains, plasmids, and primers

All other experiments were done using SK1-derived yeast strains (ho::*LYS2 lys2 ura3 leu2*::hisG *his3*::hisG *trp1*::hisG), which are listed in Table S2. The exception is the use of strains from the BY-deletion collection (Winzeler et al. 1999) that were crossed to SK1 WT cells which are described below. Split-GFP reporters consisted of pRS315-*NOP1pr-GFP*_*11*_*-mCherry-PUS1* (pSJ1321) for the INM and pRS315-*NOP1pr-GFP*_*11*_*-mCherry-SCSTM* (pSJ1568) for the ONM/ER, whose construction has been previously described in detail (Smoyer et al. 2016). Standard techniques were used for DNA and yeast manipulations, including C-terminal tagging with fluorescent proteins and gene deletion by PCR-based methods (Gardner and Jaspersen 2014). PCR primers for gene deletion were designed as follows: F1 primer-60 bp upstream of gene-specific start codon followed by *cggatccccgggttaattaa*; R1 primer-60 bp downstream of gene specific stop codon on the reverse strand followed by *tcgatgaattcgagctcgt*. PCR primers to tag genes at the C-terminus were designed as follows: F5 primer-60 bp of gene-specific sequence immediately before the stop codon followed by *ggtgacggtgctggttta*; R3 primer-60 bp of gene specific sequence immediately after the stop codon on the reverse strand followed by *tcgatgaattcgagctcg* (Gardner and Jaspersen 2014).

### Yeast growth, fixation and imaging

Sporulation was induced by starvation. Diploid yeasts were first grown for 24 h at 30°C in eSCM with agitation. Following growth in enriched media, cells were washed twice with water and twice with SPO, before resuspension in SPO and incubation at 30°C for 24-48 h with shaking. In most cases, cells were then imaged live.

To determine sporulation efficiency of heterozygotes, the haploid strain lacking the chosen gene from the BY-deletion collection (Winzeler et al. 1999) was mated to the haploid SK1 wild-type strain to generate the heterozygous deletion strains used in this study. All homozygous deletion strains were generated by PCR-mediated deletion of genes in SK1. Cells were fixed by pelleting 1 mL of the meiotic culture and resuspending the pellet in 4% paraformaldehyde solution (4.5% sucrose, 4% paraformaldehyde), followed by incubating the cells at room temperature with rotation for 15 min. Cells were washed three times with phosphate buffered saline PBS (8 mM Na_2_HPO_4_, 2 mM KH_2_PO_4_, 137 mM NaCl, 2.7 mM KCl), followed by treating the cells with 0.05 μM of DAPI in permeabilization buffer (1% TritonX-100, 0.1 M potassium phosphate, 1.2 M sorbitol). Cells were then washed with potassium/sorbitol (0.1 M potassium phosphate, 1.2 M sorbitol) and imaged. Sporulation efficiency was determined in three biological replicates by counting the number of asci with four, three, and two nuclei divided by the total number of cells.

Live cell images showing the 2:2 segregation pattern and the time-lapse images were acquired with cells immobilized in a CellASIC Onix2 microfluidic device (Millipore) perfusing with fresh SPO media at a flow rate of 5kPa on a Nikon (Garden City, NY) Eclipse Ti2 microscope equipped with a Yokogawa CSU W1 spinning disk head and a Flash 4 camera (Hamamatsu) using a Nikon 40x/1.5na water immersion Apochromat objective. Split-GFP/GFP and mCherry were excited using laser lines at 488nm and 561nm and emission filters ET525/36M and ET605/52M respectively with alternating excitation. Z stacks were acquired with a step size of 0.6 *µ*m and time series at intervals of 5 min. Imaging data was processed using NIH ImageJ software. Fluorescence channels were max projected for visualization, and time lapse data was aligned to correct for drift using a brightfield reference channel using a custom image alignment tool based on (Thevenaz et al. 1998) available for download at https://research.stowers.org/imagejplugins/.

Additional live cell images and fixed cell images were acquired on a Leica SP8 microscope with 100X, 1.4 NA oil objective. Split-GFP/GFP and mCherry were excited using laser lines at 488 nm and 561 nm, with emission photons collected by an internal Leica HyD hybrid detector with spectral windows 492 to 555 nm for split-GFP/GFP and 496 nm to 649 nm for mCherry. DAPI was excited using a 405 nm diode laser and emission photons were collected using a PMT detector with a spectral widow of 430 to 550 nm. For post-processing, images were scaled 2 × 2 with bilinear interpolation.

## Results and Discussion

### The gamete INM proteome is distinct compared to the mitotic INM proteome

Previously, we used a split-GFP strategy to assay protein access to the INM in mitotic cells (Smoyer et al. 2016). This approach to define the INM proteome takes advantage of the fact that full-length super-folder GFP can be split into two fragments between the tenth and eleventh tertiary β-barrel structure of super-folder GFP yielding two distinct fragments of GFP, GFP_1-10_ (24 kDa) and GFP_11_ (3 kDa) (Cabantous et al. 2005; Cabantous and Waldo 2006), which do not fluoresce independently; however, when the two fragments are brought together, GFP is reconstituted and fluorescent signal is detected (Figure 1A). By fusing GFP_11_ to mCherry and the gene encoding the soluble nuclear protein, Pus1, expressed under the *NOP1* promoter (*NOP1pr-GFP*_*11*_*-mCherry-PUS1*, referred to as the nuclear reporter), we could detect INM localization of proteins tagged with GFP_1-10_ at their C-terminus as fluorescent GFP signal was reconstituted at the INM (Figure 1B). A construct containing GFP_11_-mCherry fused to the transmembrane domain of Scs2 (residues 223-244, *NOP1pr-GFP*_*11*_*-mCherry-SCS2-TM*, referred to as the ONM/ER reporter) allows determination of protein localization to the ONM/ER (Figure 1B). We generated a C-terminal library considering all ORFs within the genome. Genes with an annotated transmembrane domain or predicted hydrophobic region that spanned greater than or equal to sixteen amino acids were selected to generate the library (Krogh et al. 2001; Smoyer et al. 2016). 1,063 genes were successfully fused to GFP_1-10_ at their C-terminus and were included in the library. Of these, 312 genes accessed the INM during mitotic growth, 529 were negative and the remainder were soluble (Smoyer et al. 2016). Many of the hits were bonafide INM components, however, it is important to note that a positive GFP signal in this assay does not imply that the majority of protein is present at the INM and/or protein localization is exclusive to the INM (Smoyer et al. 2016). The assay is sensitive to protein topology; the use of C-terminal tagging will overlook tail-anchored membrane proteins that have C-termini within the luminal space, for example. In mitotic cells, a N-terminal library was used to identify 100 additional proteins able to access the INM. Unfortunately, the promoter utilized in the N-terminal library is not expressed during gametogenesis so that library was not used here.

**Figure 1.**
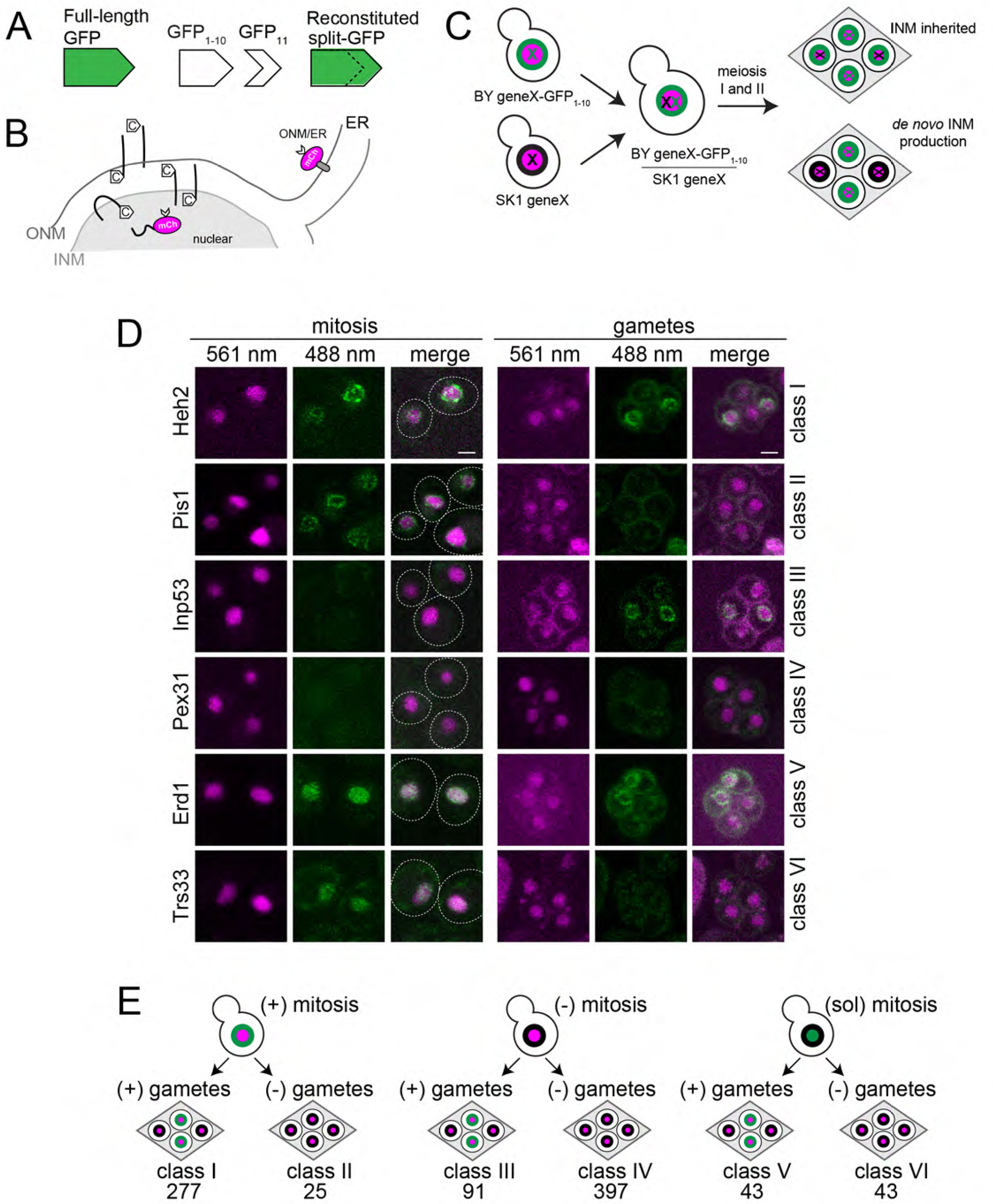
Split-GFP to define the INM proteome in gametes. (A) Schematic of the split-GFP complementation system. (B) Schematic illustrating the localization of GFP_11_-mCherry reporters used to detect signal of C-terminally GFP_1-10_ tagged proteins at the INM (GFP_11_-mCherry-Pus1) or at the ONM/ER surface (GFP_11_-mCherry-Scs2TM). (C) A heterozygous diploid strain used in the screen was generated by mating a haploid yeast containing the GFP_1-10_ tagged gene (green X) and an SK1 yeast with an untagged gene (black X). After meiosis I and II, gametes will inherit a single copy of gene X; two possibilities of protein expression at the INM in gametes are shown. (D-E) Protein expression in mitosis and meiosis can be categorized into six classes. (D) Representative images of proteins in each class showing the reconstituted GFP at 488 nm (green) and GFP_11_-mCherry-Pus1 at 561 nm (magenta). A merged image along with the cell outline is also shown. Residual background is autofluorescence, which was shown by multispectral imaging (not depicted). Bar, 2 *µ*m. (E) Summary of six classes with the number of proteins in each class shown.

To define the INM proteome in gametes, we crossed our C-terminal library to wild-type SK1 yeast (Figure 1C). This resulted in a heterozygous diploid yeast that efficiently underwent the two meiotic divisions to form four gametes (also known as spores), which are packed into tetrads. The BY strain in which the C-terminal library was constructed does not readily undergo meiosis, so mating to SK1 was necessary for efficient gamete formation (typically 50-80% tetrads after 48 h in sporulation media). Neither the SK1 strain background nor heterozygosity affected mitotic protein localization (Figure S1A), consistent with the idea that there are few proteomic changes between haploid and diploid cells during mitosis (de Godoy et al. 2008). The BY/SK1 hybrid strain also behaved similarly with respect to a SK1 diploid strain in terms of protein localization in gametes (Figure S1B). In yeast, meiosis is generally induced by nitrogen and phosphate starvation. Analysis of randomly selected clones from our library showed that protein distribution was also largely unaffected by growth in sporulation media compared to the rich media used for mitotic growth (Table S3). We screened the heterozygous diploid library for protein localization to the INM after cells were induced to undergo the meiotic program and form gametes. Although meiosis was not synchronous, in most cultures, more than 50% of cells formed tetrads, allowing us to efficiently score INM distribution in four spored meiotic progeny. Examples are shown in Figure 1D, and the results are summarized in Figure 1E and in Table S1.

The nucleus is an essential organelle, and we assumed that much of the NE and associated INM proteins would be inherited by the gametes (Figure 1C). If the INM was inherited, we expected all four meiotic products to be either positive or negative due to inheritance of synthesized proteins and the NE from the parental cell (Figure 1C). Interestingly, all of the positive results in gametes (411) showed a 2:2 segregation pattern – two nuclei contain reconstituted GFP signal at the INM and two nuclei lack signal. This pattern is consistent with *de novo* production of INM components within gametes that inherit the tagged gene. While *de novo* production was possibly expected for a few specific nucleoporins (e.g. Pom33 and Pom34) based on recent findings of NPC remodeling during meiosis (King et al. 2019), our data suggest that the bulk of the INM proteome is resynthesized in each gamete. Using heterozygous diploids that contained full-length GFP and mCherry tagged proteins, we confirmed this 2:2 segregation pattern for INM components (Figure S2). Thus, the INM is distinct in gametes compared to mitotic cells, with most, if not all, INM protein made *de novo* in gametes rather than being inherited from the parental cell.

### Protein expression at the INM is distinct from total protein expression

We classified our results by considering the individual protein localization to the INM in mitosis compared to the specific protein localization in gametes and found the following (Figure 1E): (I) signal at the INM in both mitosis and gametes; (II) signal at the INM in mitotic cells but not in gametes; (III) signal at the INM in gametes but not in mitotic cells; (IV) no INM signal in mitosis or gametes; (V) a soluble nuclear signal in mitotic cells but a signal at the INM in gametes; and (VI) a soluble nuclear signal in mitotic cells but no signal in gametes. Most genes fell into class I or class IV, suggesting that they are likely constitutive components of the INM or components of other organelles, respectively.

Twenty-five class II genes were present at the INM during mitosis but were not observed at the INM in gametes while ninety-one class III genes were observed at the INM only in gametes. Due to the limited number of proteins in class II and class III, we were not able to perform meaningful GO term analysis on either class. Manual inspection of the ninety-one genes within class III suggested they have similar or related functions that could be grouped: eleven function in cell wall biosynthesis, eleven are involved in various lipid biosynthetic and metabolic pathways, four genes function in protein degradation pathways, and twelve have unknown functions. Rtn1 and Rtn2 belong to the reticulon family of proteins that are involved in membrane curvature and NPC assembly (Dawson et al. 2009), and both were identified as class III genes.

Gametogenesis is regulated by a cascade of transcription factors (Chu et al. 1998; Neiman 2011; Xu et al. 1995). It seemed likely that changes in INM localization in class II and class III could simply be attributed to differences in protein expression during vegetative growth and meiosis. However, only two genes in class III (*DTR1* and *RTN2*) are reported to be transcriptionally regulated by Ndt80, the master regulator of meiotic gene expression (Chu et al. 1998). To further evaluate the role that transcription and translation might play in our results, we compared protein expression levels of individual genes in class II and class III in mitotic cells to gametes using published total protein amounts determined throughout meiosis using quantitative mass spectroscopy (Cheng et al. 2018). We did not observe a general pattern where proteins showed higher expression in mitotic cells compared to gametes as we expected for class II (Figure 2A) nor did we observe an overall lower expression pattern in mitotic cells compared to gametes as expected for class III (Figure 2B). Rtn2 was a significant exception; as a target of Ndt80 (Chu and Herskowitz 1998), its transcription and translation are both upregulated in meiosis compared to mitosis. Total protein levels of the other yeast reticulon, Rtn1, indicate that it is significantly more abundant in mitotic cells compared to meiotic cells, although we only detected it at the INM in gametes (class III).

**Figure 2.**
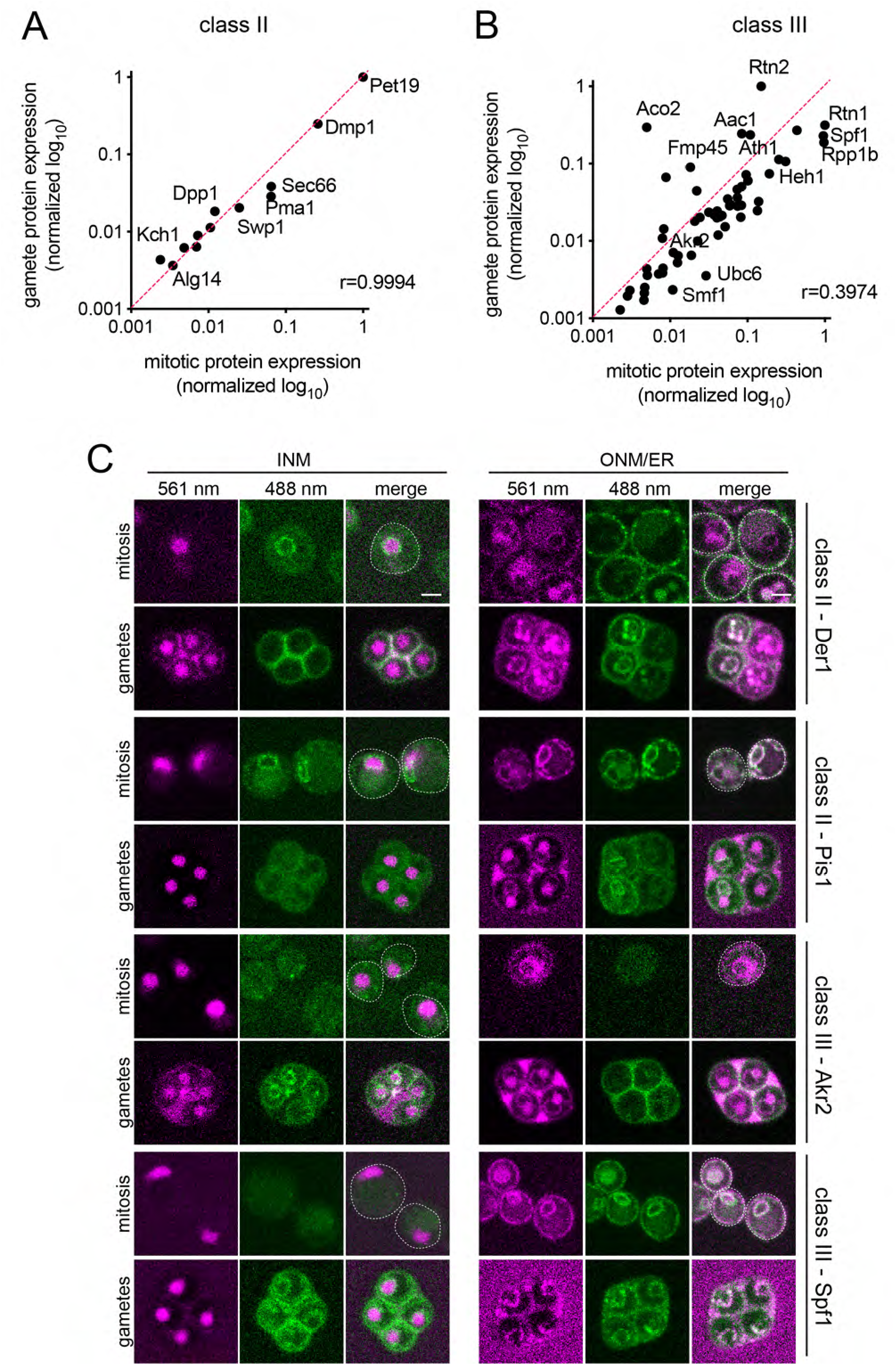
Total cellular protein compared to protein specifically at the INM. (A-B) Total protein in mitotic cells plotted against total protein in gametes for proteins in class II (A) and class III (B) using data from (Cheng et al. 2018). The line through the origin with a slope of 1 is for ease of viewing and would be expected for a perfect correlation. The Pearson correlation coefficient is included on graph. Protein levels were normalized within each class with the highest value in class II (A) or class III (B) set to 1 and all other values normalized to that value. (C) Protein localization at the INM (using GFP_11_-mCherry-Pus1) or the ONM/ER (using GFP_11_-mCherry-Scs2TM) was assayed in mitotically growing cells or gametes for the indicated proteins. Fluorescence at 561 nm (magenta) and 488 nm (green) is shown along with a merged image that contains the cell outline. Residual background is autofluorescence, which was shown by multispectral imaging (not depicted). Bar, 2 *µ*m.

Next, we considered the possibility that proteins in class II might be expressed in gametes but not localize to the INM. Similarly, class III proteins may be expressed in mitotic cells but not localize to the INM. To test this idea, we used the ONM/ER reporter construct designed to detect protein localization to the ONM and cortical ER. We found that Der1, a class II protein, localized to the INM, as expected, and the cortical ER in mitosis (Figure 2C). In gametes, we confirmed that Der1 is not present at the INM; however, it is present at the ONM and cortical ER (Figure 2C). The class II protein Pis1 shows a similar distribution to Der1, except it is also detected at both the cortical and perinuclear ER (Figure 2C). The class III protein, Akr2, was absent at the INM in mitotic cells and present in gametes, as expected, and it did not localize to the ONM/ER in mitotic cells or gametes (Figure 2D). Another class III protein, Spf1, was visualized at the perinuclear and cortical ER in mitotic cells and gametes (Figure 2D). Class I and IV proteins can also be detected with the ONM/ER reporter in mitotic cells and gametes (Figure S3).

Analysis of unassembled subunits of protein complexes in mitotic cells showed they accumulate at the INM for turnover by quality control pathways involving the Asi1 ubiquitin ligase (Natarajan et al. 2020). It is unknown if this Asi1-mediated pathway is functional in gametes, but our screen showed that Asi1 and its paralog Asi3 are present at the INM in gametes as well as mitotic cells (Table S1), raising the possibility that protein quality control may underlie the localization patterns we observe in class II and class III. Consistent with this idea, four of the twenty-five proteins in class II are Asi1 targets: Dpm1, Pmt3, Der1, and Pma1 (Khmelinskii et al. 2014; Smoyer et al. 2019). However, class III does not contain any known Asi1 targets, although it does include several E2 ligases and other factors implicated in protein quality control. It is therefore difficult to understand how Asi1-mediated degradation alone leads to the distribution patterns we observe. In fact, multiple Asi1 targets are found in class I, which does not show changes in protein distribution.

Taken together, these data suggest that the pattern of protein localization we observe at the INM in gametes is not due to gene activation/repression by the meiotic transcriptional cascade. In addition, protein quality control is unlikely to contribute to INM distribution beyond a handful of targets in class II. We cannot determine from our current data the overall role that protein degradation plays in sculpting the INM proteome given that other ligases and/or Asi1/Asi3 targets could be involved in meiosis. Below, we present additional analysis aimed at understanding the role class III genes play at the INM and targeting mechanisms.

### Class III genes localized to the INM are needed for gamete formation

We hypothesized that the INM distribution of at least some class III genes are important to meiotic and/or gamete specific INM function. To further investigate the functional relevance of proteins in class III, we evaluated the effect gene deletions had on gamete formation. If the protein performs an essential function at the INM in gametes, then we anticipated that a partial loss of gene function would lead to a reduction in gamete formation. Heterozygous diploids containing individual gene deletions were made by crossing strains from the BY yeast deletion collection (Winzeler et al. 1999) with wild-type SK1 cells. After induction of sporulation for 48 h, gamete formation was assayed microscopically to determine the fraction of cells that developed into well-formed asci containing the four meiotic products (tetrads), known as the sporulation frequency. Compared to wild-type cells, 41 and 25 of the 76 heterozygous deletion strains had a mild or moderate decrease in gamete formation, defined as greater than 2-fold or 4-fold reduction in sporulation frequency, respectively (Table S4). Heterozygous deletions of four class III genes resulted in a 10-fold reduction in gamete formation compared to wild-type: *FMP45*, an integral membrane protein involved in sphingolipid biosynthesis (Young et al. 2002); *TLG2*, a syntaxin-like t-SNARE (Bryant and James 2001); *KTR7*, a putative mannosyltransferase (Lussier et al. 1997); and *SRC1/HEH1*, a LEM-domain containing protein (Grund et al. 2008; Rodriguez-Navarro et al. 2002). The *FMP45* paralog, *YNL194c*, is also a class III gene with moderate defects in sporulation frequency, consistent with previous work showing a role for sphingolipid synthesis in gamete formation (Young et al. 2002). Another set of paralogs, the yeast reticulons (Oertle et al. 2003), were amongst the class III genes with mild (*RTN2*) or moderate (*RTN1*) sporulation defects. The most severe defect in gamete formation observed in heterozygous diploids was in *heh1Δ/HEH1* with a mean sporulation frequency of 3.4±0.9% compared to 68.2±3.3% in wild-type cells.

To further investigate a role for *HEH1* and the reticulons in gamete formation, we created homozygous deletion strains in the SK1 background. Perhaps not unexpectedly, we observed defects in gamete formation that increased in severity in homozygous deletions where gene function was completely lost (Figure 3). Analysis of nuclear morphology using DAPI suggests that the defect in gamete formation is not caused by a failure to enter meiosis as dyads are observed in addition to tetrads in the deletion strains (Figure 3). However, the fact that few mature gametes with four nuclei were recovered points to a role for Heh1, Rtn1 and Rtn2 in meiotic progression and/or gamete formation. It is likely that these defects stem at least in part from gene function at the INM, however, future experiments will be needed to determine their specific INM role in gamete maturation.

**Figure 3.**
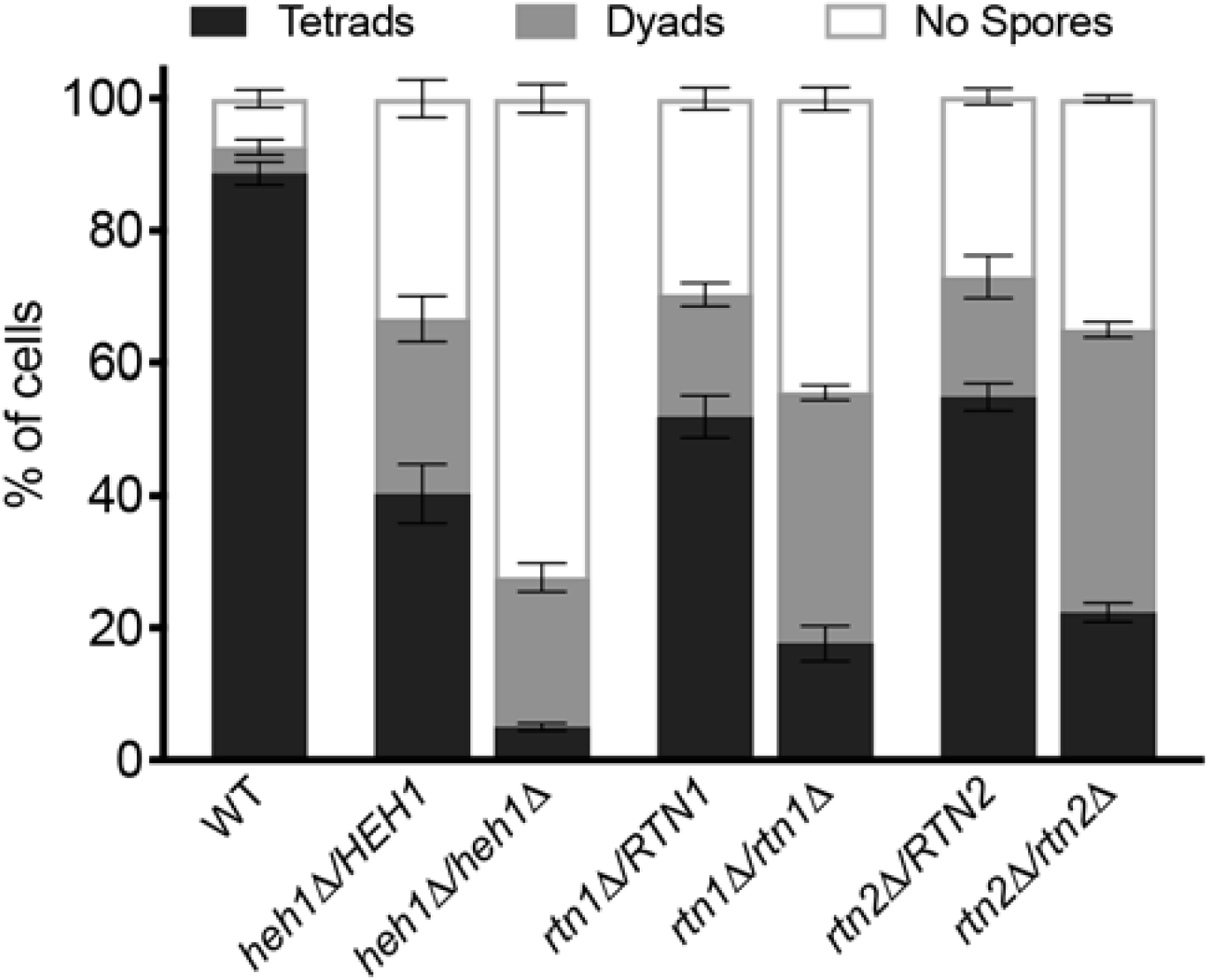
Class III proteins are necessary for gamete formation late in meiosis II. Sporulation of wild-type and heterozygous or homozygous deletions of *HEH1, RTN1* and *RTN2*. The average fraction of cells that formed an ascus with four nuclei (tetrad), or two nuclei (dyad) compared to the number of cells that did not sporulate. 100 cells from three biological replicates were tested. Error bars, SEM. Unpaired t-test was used for statistical analysis, with all mutants having a p<0.001 compared with wild-type.

### Large proteins access the INM in gametes

In budding yeast, the NE remains intact throughout the lifecycle (Moens 1971; Moens and Rapport 1971). Two mechanisms are thought to facilitate protein targeting to the INM in mitotic cells. Proteins with small extraluminal domains can use a diffusion-retention mediated pathway similar to that of small soluble proteins (Gorlich and Kutay 1999; Katta et al. 2014). Diffusion occurs through peripheral NPC channels and the protein is maintained at the INM through interactions with chromatin or other nuclear proteins (Figure 4A). For larger proteins, a transport-mediated pathway is necessary. A nuclear targeting sequence is recognized by karyopherins that bind the protein and mediate Ran-GTP-dependent transport through the NPCs (Figure 4A).

**Figure 4.**
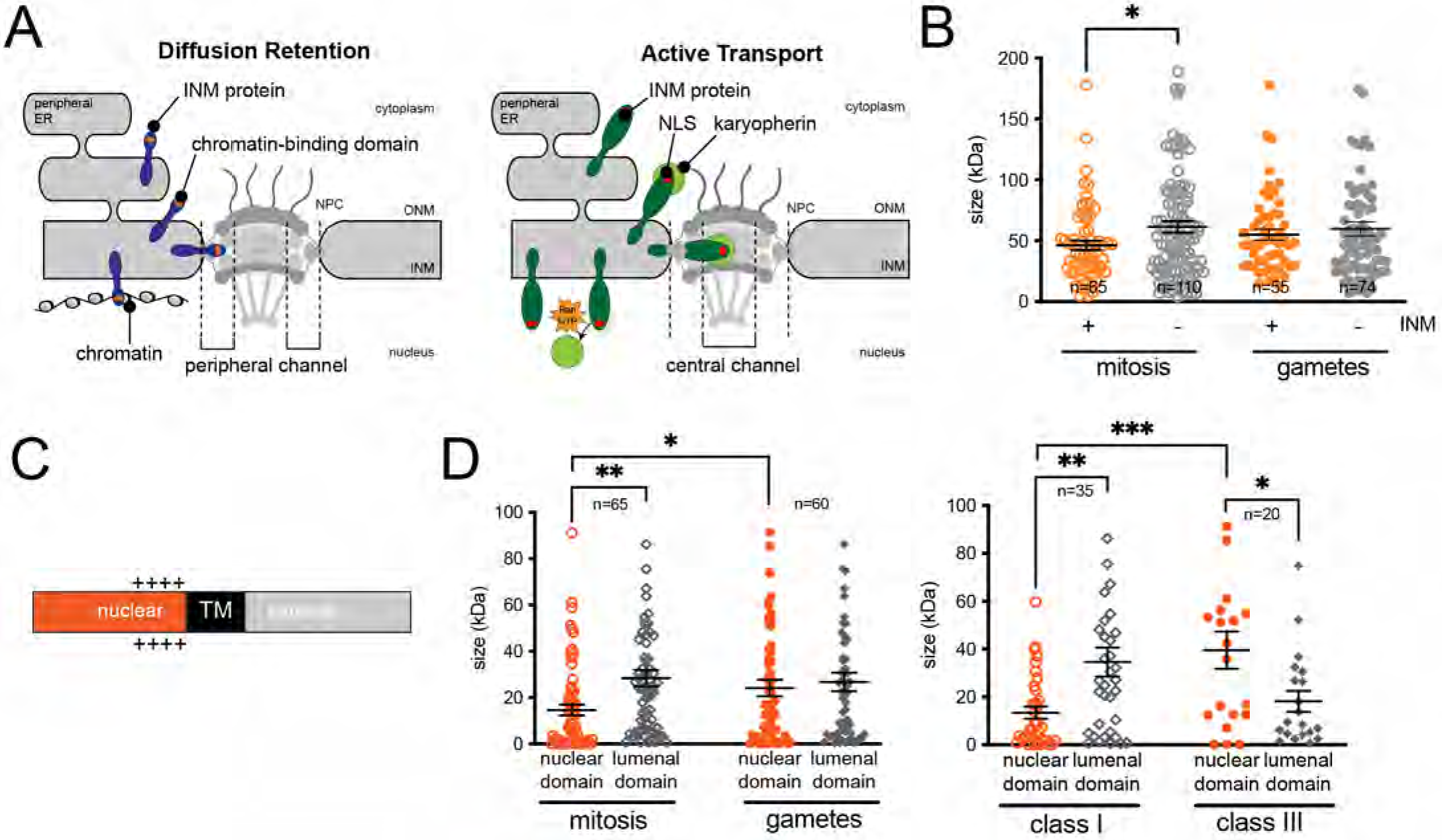
High molecular weight proteins access the INM in gametes. (A) Schematic showing the two mechanisms utilized by INM proteins to localize to the nucleus following their synthesis on the ER membrane. Adapted from Katta et al. 2014. (B) The molecular weight of proteins that localize to the INM (+) compared to proteins that do not localize to the INM (-) was compared for mitotic cells and gametes. P-values were calculated using the Student’s t-test. All were not significant, except *, p=0.03. n, indicated. Error bars, SEM. (C) Schematic representation of a single pass transmembrane domain protein showing the positive charges that are on the nuclear side of the protein (Smoyer et al. 2016). (D) The molecular weight of the nuclear domain and luminal domain of single pass transmembrane proteins that target the INM during mitosis were compared to gametes (left) and class I (INM mitosis and gametes) was compared to class III (INM gametes only). P-values were calculated using the Student’s t-test. Statistically significant values are indicated: *, p<0.05; **, p<0.001; ***, p<0.0001. n, indicated. Error bars, SEM.

No particular motif was enriched above background in class I and class III, the two groups with protein present at the INM in gametes. This suggests that a simple localization sequence is unlikely to confer INM localization in gametes, similar to mitotic cells (Smoyer et al. 2016). Previously, we showed that proteins destined for the INM in mitotic cells show a significantly lower total molecular weight compared to proteins unable to reach the INM (Smoyer et al. 2016). Interestingly, this size preference for INM proteins is not maintained in gametes (Figure 4B; Table S5), perhaps indicating a change in the diffusion barrier. Examination of the molecular mass of single-pass transmembrane proteins (Figure 4C) showed that the strong preference for proteins with small extraluminal domains in mitotic cells was lost in gametes (Figure 4D). This is most evident in a comparison of class I (at the INM in both mitosis and gametes) and class III (present only in gametes): the average extraluminal domain in class I is 13.4±2.6 kDa (n=35) compared to 39.6±7.8 kDa (n=20) for class III. These findings support the notion that there is a loss and/or change in the nuclear permeability barrier during a late stage in gametogenesis that allows larger proteins and proteins with large nuclear domains to access the INM in gametes.

### Transient loss of the nuclear permeability barrier after anaphase II

To test the idea that NE permeability is altered during gametogenesis, we examined NE integrity using the soluble nuclear reporter, Pus1-GFP. Cells were followed through meiosis I and meiosis II until spores formed, using nuclear morphology and the transmitted light image to stage the cells. At the end of anaphase II, Pus1-GFP was present in nuclei and in the fifth compartment, but shortly after anaphase the nuclear signal dissipated throughout the cell. A soluble nuclear signal eventually returned in four distinct nuclei (Figure 5A). The transient loss of the soluble signal was not unique to Pus1-GFP but was also observed for the nuclear protein Lys21-GFP (Figure 5B) (Fuchs and Loidl 2004). A non-soluble protein that is attached to chromatin, histone H2B, remains associated with DNA throughout meiosis and spore formation, including time points when both soluble proteins become dispersed (Figure 5A-C). EM data shows an intact NE throughout meiosis (King et al. 2019), however, our data suggests a loss of the diffusion barrier occurs at least for soluble proteins.

**Figure 5.**
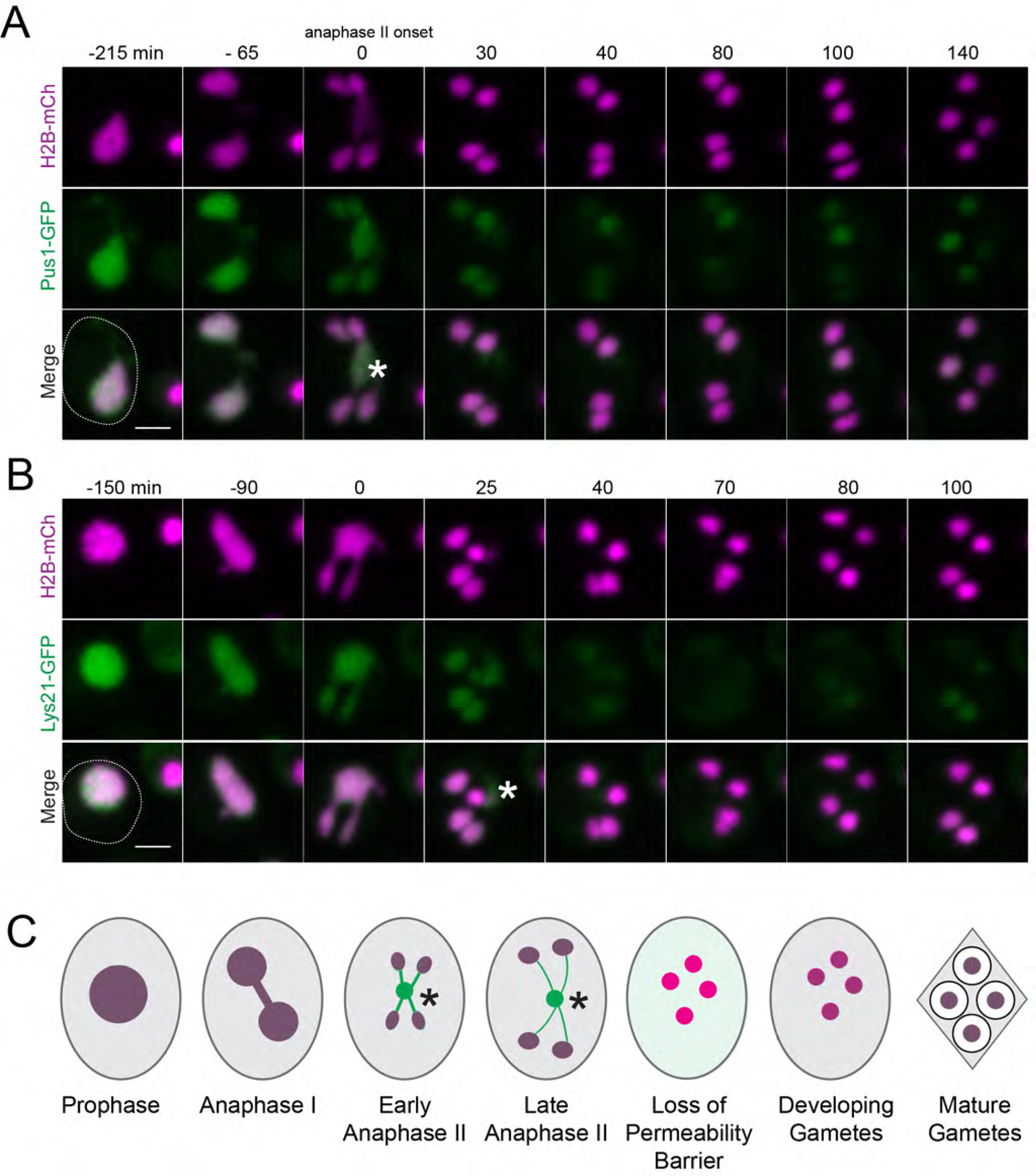
Transient loss of the nuclear permeability barrier for soluble proteins. (A) Montage of a cell containing Pus1-GFP (green) and Htb2-mCherry (magenta) progressing through meiosis. (B) Montage of a cell containing Lys21-GFP (green) and Htb2-mCherry (magenta) progressing through meiosis as in (A). The onset of anaphase II was defined as t=0 min. Asterisk marks the location of the fifth compartment; the cell outline is depicted. Bar, 2 *µ*m. (C) Schematic of transient loss of NE compartmentalization based on data from (A-B), with green and magenta representing the soluble and chromatin-associated protein, respectively. An asterisks marks the location of the fifth compartment.

### Conclusion

We show that the INM proteome in gametes is distinct from mitotic cells and that the expression of proteins in gametes is important for gamete formation. The 2:2 segregation pattern observed for all INM components strongly suggests that gametes make INM proteins *de novo* rather than inheriting these proteins from the parental cell. We hypothesize that most, if not all, proteins at the INM are retained in the fifth compartment and eliminated as debris through vacuolar lysis during gamete maturation, similar to the pathway used to clear most nucleoporins (King et al. 2019; King and Unal 2020).

Surprisingly, the preference of smaller proteins and smaller extraluminal domains was lost in gametes compared to mitotic cells, indicating that gametes undergo a change in the nuclear permeability barrier. In fission yeast, the NE diffusion barrier is compromised during early anaphase B of meiosis II in a phenomenon known as virtual NE breakdown (vNEBD) (Arai et al. 2010; Asakawa et al. 2011). During vNEBD, NPCs remain intact, but a change in localization of the RanGAP1 from the cytoplasm to the nucleus leads to a collapse of the Ran-GTP gradient. As a result, nuclear and cytoplasmic contents to undergo mixing (Asakawa et al. 2016). Breakdown of the NE or loss of the diffusion barrier has not been reported in budding yeast at any stage of its lifecycle (Boettcher and Barral 2013). However, we observed a transient loss of the nuclear permeability barrier for soluble proteins. It is unknown if the appearance of class III proteins (or disappearance of class II proteins) at the INM correlates with this change in permeability, with NPC remodeling or other events that alter the diffusion barrier, including vNEBD or possibly vacuolar lysis, which changes intracellular pH and releases multiple proteases.

## Acknowledgements

We thank Sarah Zanders, Gloria Brar and Elçin Ünal for strains and are grateful to Sarah Zanders, Grant King and the Jaspersen lab for helpful suggestions throughout the project and for their comments on the manuscript. Original data underlying this manuscript can be downloaded from the Stowers Original Data Repository at http://www.stowers.org/research/publications/libpb-1637. Research reported in this publication was supported by the Stowers Institute for Medical Research. The funder had no role in study design, data collection and analysis, decision to publish, or preparation of the manuscript. The authors declare no competing financial interests. Strains and plasmids are available upon request. The authors affirm that all data necessary for confirming the conclusions of the article are present within the article, figures, and tables.

## Author contributions

SLJ and SNS conceived the experiments, SNS constructed strains, SNS and SES performed the screen, and SNS did the follow-up analysis with help from SES in image acquisition and SES and JRU in data analysis. SNS and SLJ prepared figures and wrote the paper with input from all the authors.

## Supplementary Information

**Figure S1.**
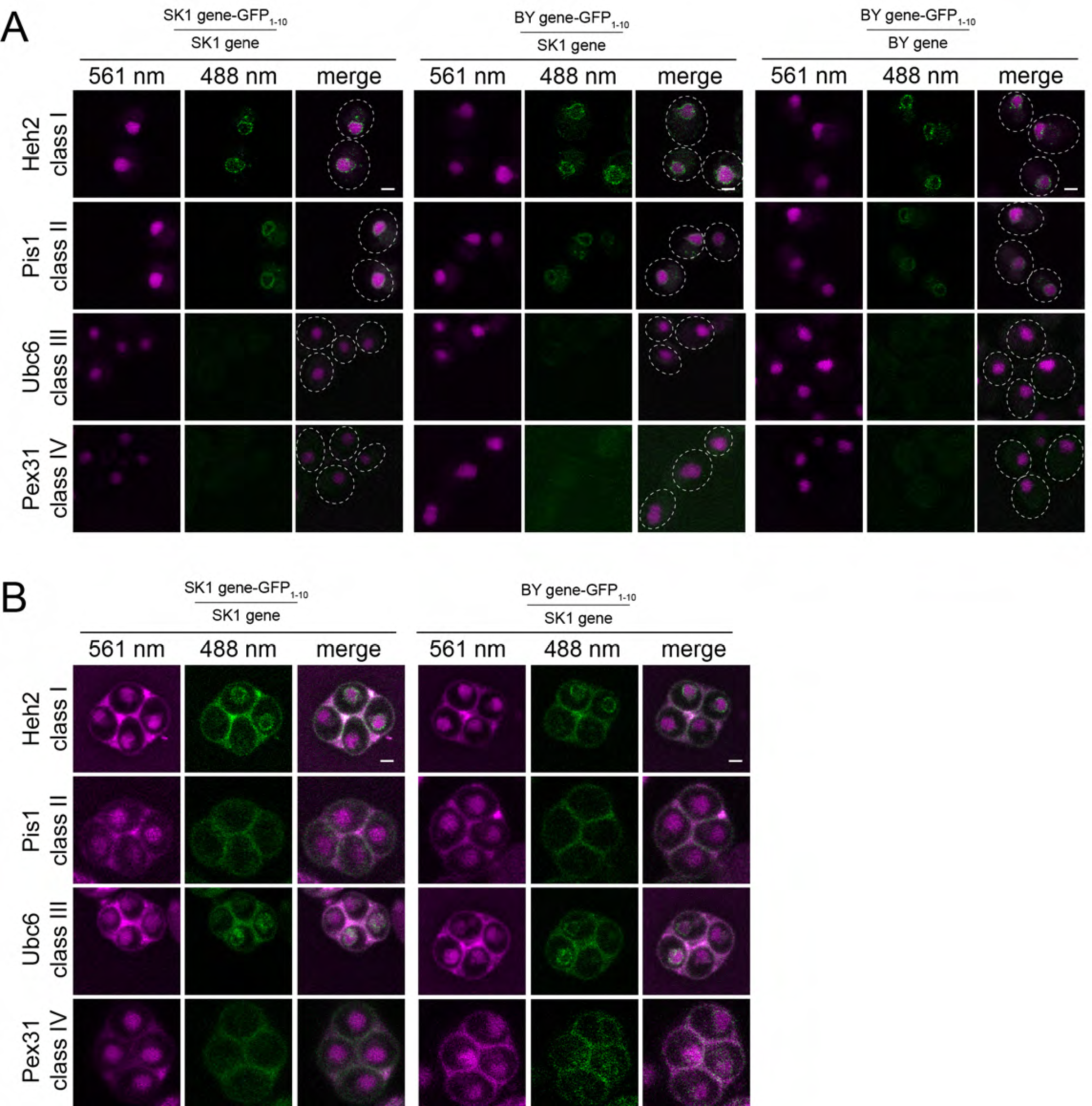
INM protein localization is not strain specific. (A) Mitotic localization of proteins from class I-IV in three different diploid strains with the indicated genotype. (B) INM localization of a representative class I-IV protein in gametes derived from two different diploid strains. Reconstituted GFP is at 488 nm (green) and GFP_11_-mCherry-Pus1 is at 561 nm (magenta). A merged image along with the cell outline is also shown. Residual background is autofluorescence, which was shown by multispectral imaging (not depicted). Bar, 2 *µ*m.

**Figure S2.**
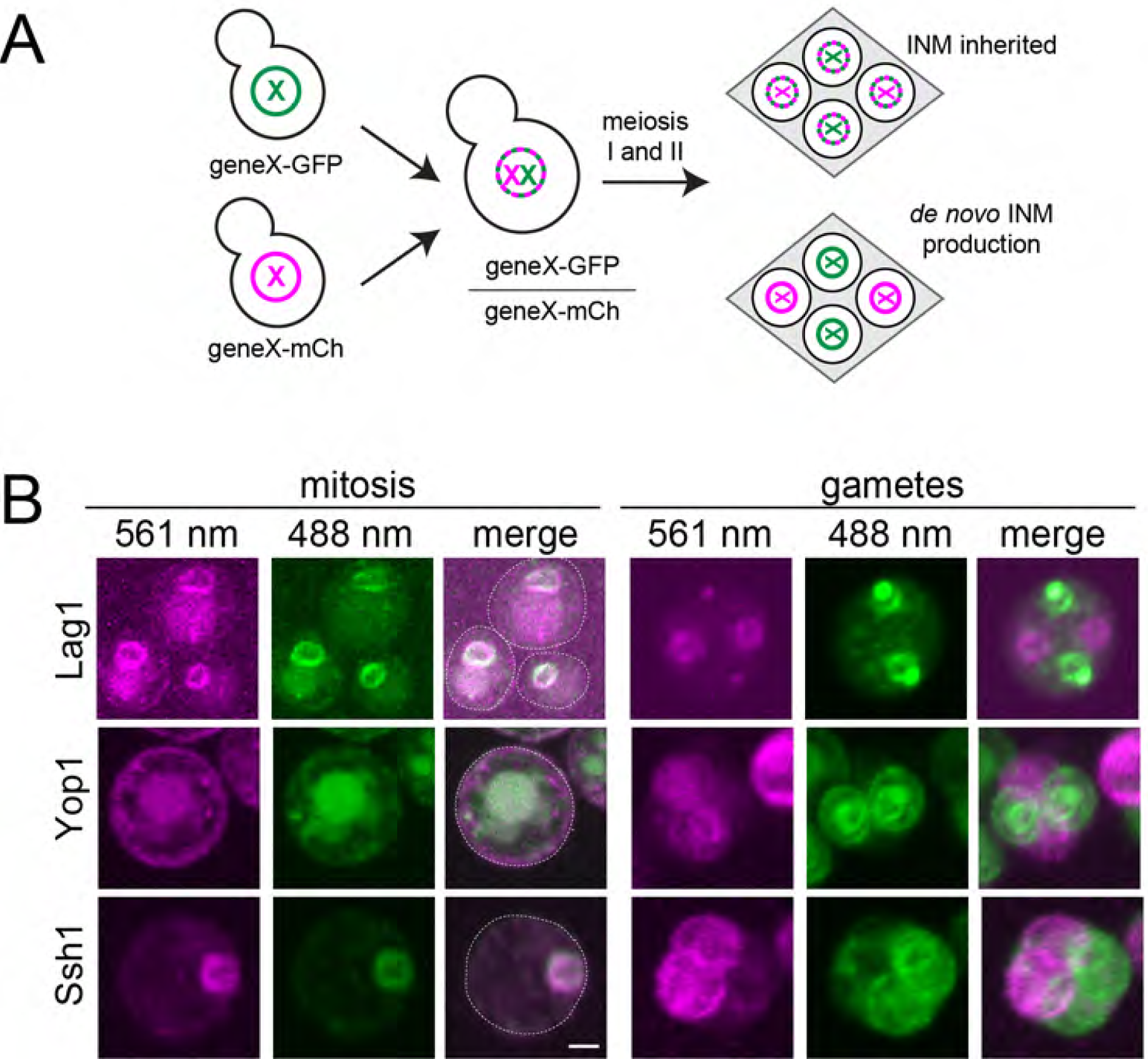
The INM proteome is made *de novo* in gametes. (A) In a heterozygous diploid strain in which one copy of the gene was tagged with GFP (green X) and the other copy was tagged with mCherry (magenta X), gametes will inherit a single copy of each tagged gene. Two possibilities of protein expression in gametes are shown. (B) Representative images from heterozygous diploids showing protein distribution in mitotic cells and in the four gametes. A merged image along with the cell outline for mitotic cells is also shown. Residual background is autofluorescence, which was shown by multispectral imaging (not depicted). Bar, 2 *µ*m.

**Figure S3.**
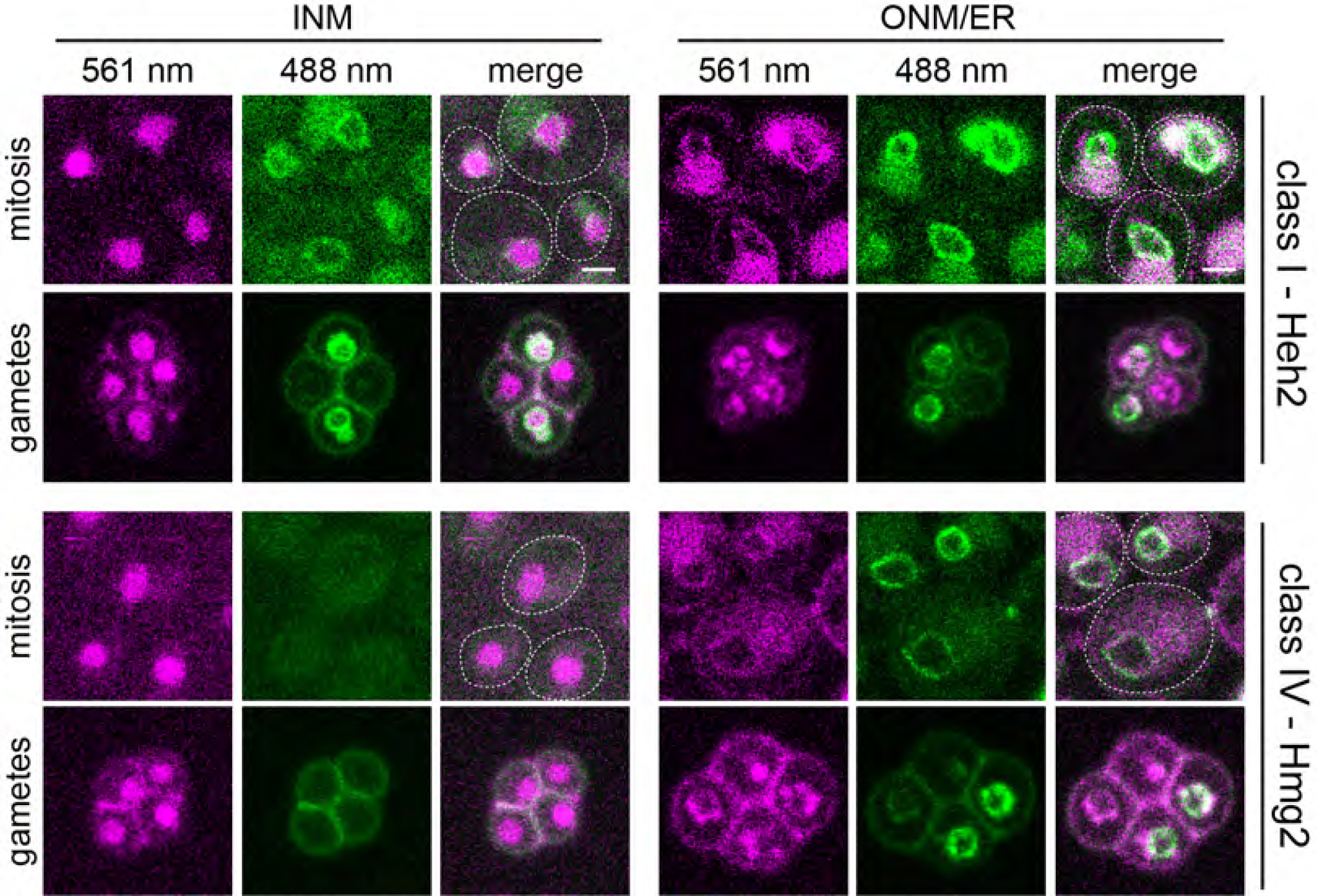
Class I and class IV proteins can be present at the ONM/ER. Protein localization at the INM (using GFP_11_-mCherry-Pus1) or the ONM/ER (using s GFP_11_-mCherry-Scs2TM) was assayed in mitotically growing cells or gametes for the indicated proteins. Fluorescence at 561 nm (magenta) and 488 nm (green) is shown along with a merged image that contains the cell outline. Residual background is autofluorescence, which was shown by multispectral imaging (not depicted). Bar, 2 *µ*m.

